# GATA1-deficient human pluripotent stem cells generate neutrophils with improved antifungal immunity that is mediated by the integrin CD18

**DOI:** 10.1101/2024.10.11.617742

**Authors:** Andrew S. Wagner, Frances M. Smith, David A. Bennin, James A. Votava, Rupsa Datta, Morgan A. Giese, Wenxuan Zhao, Melissa C. Skala, Jing Fan, Nancy P. Keller, Anna Huttenlocher

## Abstract

Neutrophils are critical for host defense against fungi. However, the short life span and lack of genetic tractability of primary human neutrophils has limited *in vitro* analysis of neutrophil-fungal interactions. Human induced pluripotent stem cell (iPSC)-derived neutrophils (iNeutrophils) are a genetically tractable alternative to primary human neutrophils. Here, we show that deletion of the transcription factor GATA1 from human iPSCs results in iNeutrophils with improved antifungal activity against *Aspergillus fumigatus*. GATA1 knockout (KO) iNeutrophils have increased maturation, antifungal pattern recognition receptor expression and more readily execute neutrophil effector functions compared to wild-type iNeutrophils. iNeutrophils also show a shift in their metabolism following stimulation with fungal β-glucan, including an upregulation of the pentose phosphate pathway (PPP), similar to primary human neutrophils *in vitro*. Furthermore, we show that deletion of the integrin CD18 attenuates the ability of GATA1-KO iNeutrophils to kill *A. fumigatus* but is not necessary for the upregulation of PPP. Collectively, these findings support iNeutrophils as a robust system to study human neutrophil antifungal immunity and has identified specific roles for CD18 in the defense response.

**Author Summary:** Neutrophils are important first responders to fungal infections, and understanding their antifungal functions is essential to better elucidating disease dynamics. Primary human neutrophils are short lived and do not permit genetic manipulation, limiting their use to study neutrophil-fungal interactions *in vitro*. Human induced pluripotent stem cell (iPSC)-derived neutrophils (iNeutrophils) are a genetically tractable alternative to primary human neutrophils for *in vitro* analyses. In this report we show that GATA1-deficient iPSCs generate neutrophils (iNeutrophils) that are more mature than wild-type iNeutrophils and display increased antifungal activity against the human fungal pathogen *Aspergillus fumigatus*. We also show that GATA1-deficient iNeutrophils have increased expression of antifungal receptors than wild-type cells and shift their metabolism and execute neutrophil antifungal functions at levels comparable to primary human neutrophils. Deletion of the integrin CD18 blocks the ability of GATA1-deficient iNeutrophils to kill and control the growth of *A. fumigatus*, demonstrating an important role for this integrin in iNeutrophil antifungal activity. Collectively, these findings support the use of iNeutrophils as a model to study neutrophil antifungal immunity.

## Introduction

Invasive fungal infections (IFIs) are a significant health problem with greater than 1.5 million annual deaths worldwide [1]. Invasive aspergillosis (IA), primarily caused by *Aspergillus fumigatus,* is amongst the most common IFIs that requires hospitalization, and accounts for an estimated economic burden of 1.8 billion dollars within the United States alone [2]. Patients with neutropenia are amongst the most at risk for developing IA, since neutrophils act as important first responders at early stages of disease development [3]. Consequently, understanding how neutrophils interact with *A. fumigatus* is essential to elucidate the mechanisms of disease progression.

Neutrophils are key responders to *A. fumigatus*, and as such express a repertoire of antifungal pattern recognition receptors (PRRs) that facilitate their interactions with invading fungi [4, 5]. Once engaged, PRR activation stimulates neutrophils to perform numerous effector functions. These include antimicrobial activities like phagocytosis, reactive oxygen species (ROS) production, neutrophil extracellular trap (NET) extrusion, degranulation and the release of antimicrobial peptides such as elastase and lactoferrin to kill or attenuate fungal growth [6–10]. As part of their effector functions neutrophils rapidly produce ROS to mediate pathogen control [11–13], and recent evidence has demonstrated that metabolic reprogramming following activation towards the pentose phosphate pathway is necessary for the generation of more ROS during an oxidative burst [14, 15]. Inhibiting the pentose pathway has also been shown to attenuate the antifungal activity against *A. fumigatus in vitro*, thus highlighting the importance of metabolic rewiring to control fungal growth [14]. Yet, our understanding of how human neutrophils actively sense fungal pathogens to direct metabolic changes, and how this in turn mediates their fungal killing capacity is incomplete. Primary neutrophils are short lived and are not genetically tractable [16], limiting the study of human neutrophil metabolism and host defense. Therefore, improved *in vitro* systems are needed to more effectively study fungal-neutrophil interaction dynamics.

Primary neutrophils from murine models harboring mutations of interest have been used to understand fungal defense responses. However, differences in murine granule composition, antimicrobial peptide production and overall antifungal activity have been reported [17]. Additionally, species-specific differences in the mechanisms used by neutrophils to mediate antifungal activity has complicated the translation of these findings to human neutrophils. For example, the ß(1,3)-glucan receptor dectin-1 is important in murine neutrophil antifungal activity [18, 19], but reports suggest it is dispensable for human neutrophils [10, 20]. These discrepancies highlight the need for genetically tractable human cell lines. Human myeloid cell lines such as HL- 60 and PLB-985 cells undergo neutrophil-like differentiation *in vitro* and have also been used to study neutrophil antifungal immunity. Indeed, differentiated PLB-985 cells have recently been shown to phagocytose *A. fumigatus* conidia *in vitro* [21]. However, although PLB-985 and HL-60 cells can perform antimicrobial effector functions, their capacity to do so is attenuated with reduced phagocytosis efficiency and NET release compared to primary human neutrophils [21, 22]. Human induced pluripotent stem cell (iPSC)-derived neutrophils (iNeutrophils) are a newer source of genetically tractable neutrophils that can serve as an alternative to primary human neutrophils for functional assays. Indeed, these cells have already proven effective for studying mechanisms of neutrophil migration *in vitro* [23, 24]. However, their application as a model cell line to study antimicrobial activity has been hampered by impaired functional capacity, at least in part due to heterogeneity following maturation to iNeutrophils [25–27]. In light of this, it was recently discovered that deletion of the transcription factor *GATA1*, which drives basophil and eosinophil differentiation, in iPSCs results in a more homogenous culture of mature iNeutrophils that display increased formation of NETs [28].

Here, we show that GATA1-KO iNeutrophils have increased expression of antifungal PRRs and improved antifungal activity against *A. fumigatus* compared to wild-type (WT) cells. Like primary human neutrophils, GATA1-KO iNeutrophils shift their metabolism towards the pentose phosphate pathway following stimulation with ß-glucan rich bioparticles and show a reduced redox state by single cell metabolic imaging compared to WT cells. To identify mechanisms that regulate this metabolic shift and antifungal activity, we deleted the integrin receptor CD18, an integral subunit of the ß-glucan binding protein complement receptor 3 (CR3). CD18-deficient GATA1-KO iNeutrophils had impaired ability to kill and control the growth of *A. fumigatus*. However, CD18 was not necessary for the shift in metabolism induced by ß-glucan. Collectively, this work highlights the power of iNeutrophils to dissect mechanisms that regulate human neutrophil host defense and metabolism.

## Results

### GATA1-KO iNeutrophils have increased antifungal activity against *Aspergillus fumigatus*

Given the increased maturity of GATA1-KO iNeutrophils [28], we hypothesized that these cells would display increased antifungal activity *in vitro*. To test this, we generated a new GATA1-KO iPSC line (S1 Fig) and differentiated these cells to iNeutrophils to compare the antifungal activity of WT and KO iNeutrophils against the common human fungal pathogen *A. fumigatus* strain CEA10. This fungal isolate expresses a cytoplasmic red fluorescence protein (RFP), which was used to track fungal viability (as indicated by the presence/absence of RFP signal) and growth following co-incubation with either iNeutrophils or primary human neutrophils (Fig. 1A) [29]. To assess the impact that opsonization has on antifungal activity, assays were done in media supplemented with either 2% fetal bovine serum (FBS) or 2% human serum. By 4 hours post-incubation (hpi), GATA1-KO iNeutrophils showed increased fungal clearance in both FBS and human serum treated conditions (Fig. 1B). This difference in fungal killing was most apparent in human serum supplemented media, where the mean fungal killing at 4hpi of WT iNeutrophils was ∼13% as opposed to ∼40% of fungal cells killed in GATA1-KO iNeutrophil treated conditions. Yet, it is important to note that by 4hpi, GATA1-KO iNeutrophils were still significantly less effective at clearing *A. fumigatus* germlings than primary human neutrophils. However, fungal growth at 24hpi revealed that GATA1-KO iNeutrophils attenuated *A. fumigatus* hyphal development at levels comparable to primary human neutrophils (Fig. 1C). These findings suggest that the rate at which fungal killing occurs is likely slower for GATA1- KO iNeutrophils compared to primary neutrophils. Indeed, time lapse microscopy revealed a significant delay in fungal killing between primary human neutrophils and GATA1-KO iNeutrophils over a 12-hour incubation period (Fig. 1D and S1 Movie). Nonetheless, GATA1-KO iNeutrophils still killed ∼70% of all fungal cells tracked, thus demonstrating potent antifungal activity *in vitro*.

**Figure 1:**
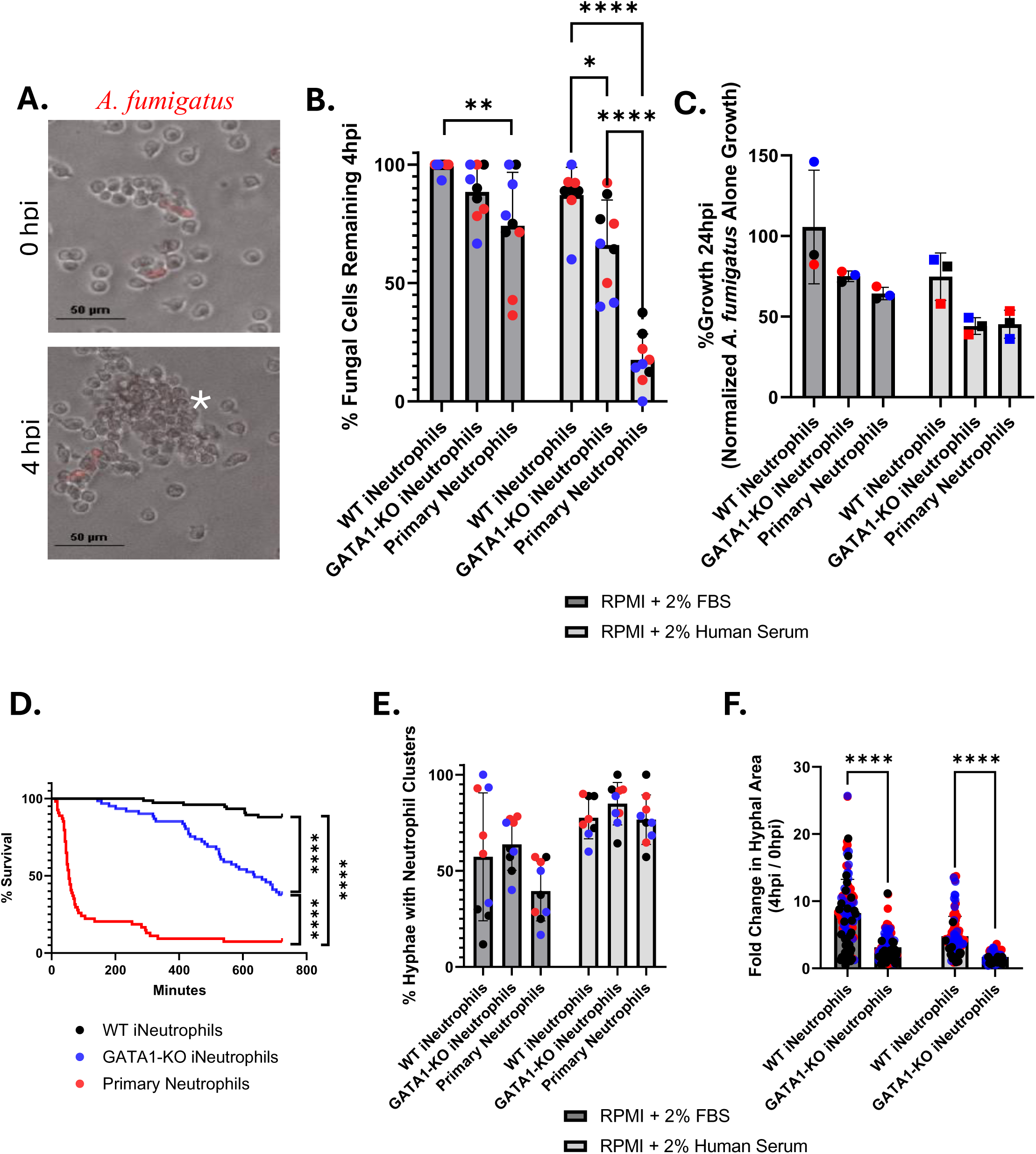
GATA1-KO iNeutrophils effectively kill and attenuate the growth of *A. fumigatus.* (A) Representative image of GATA1-KO iNeutrophils aggregating around and killing RFP expressing *A. fumigatus* germlings following 4 hours of coincubation. (White asterisks denotes fungal killing as determined by loss of cytosolic *A. fumigatus* RFP signal)(Scale bar = 50μm). (B) Quantification of *A. fumigatus* survival at 4 hpi with either iNeutrophils or primary human neutrophils in RPMI media supplemented with 2% FBS or 2% human serum. Three biological replicates (indicated by different colors) with three technical replicates were assessed for all conditions tested. (N=130 fungal cells analyzed for WT + 2% FBS, N=127 fungal cells analyzed for WT + 2% human serum, N=120 fungal cells analyzed for GATA1-KO + 2% FBS, N=94 fungal cells analyzed for GATA1-KO + 2% human serum, N=81 fungal cells analyzed for primary neutrophils + 2% FBS, N=101 fungal cells analyzed for primary neutrophils + 2% human serum)(*p<0.05, **p<0.005, ****p<0.0001, via a two-way ANOVA with Sidak’s multiple comparison test) (C) Fungal growth determined via PrestoBlue viability staining following 24 hours of coincubation with either iNeutrophils or primary human neutrophils in RPMI media supplemented with 2% FBS or 2% human serum. Growth is normalized to *A. fumigatus* samples that were not treated with (i)Neutrophils in each condition. (N=3 biological replicates)(Statistics determined via a two-way ANOVA with Sidak’s multiple comparison test). (D) Kaplan Meier survival curve of *A. fumigatus* following 12 hours of incubation with either iNeutrophils or primary human neutrophils in RPMI media supplemented with 2% human serum. Three biological replicates were run for all experimental conditions. (N=75 fungal cells tracked for WT iNeutrophils, N=61 fungal cells tracked for GATA1-KO iNeutrophils and N=57 fungal cells tracked for primary human neutrophils)(****p<0.0001 via Cox proportional hazard regression analysis). (E) Quantification of *A. fumigatus* germlings displaying clusters of five or more (i)neutrophils aggregated around them at 4 hpi in RPMI media supplemented with 2% FBS or 2% human serum. Three biological replicates (indicated by different colors) with three technical replicates were assessed for all conditions tested. (N=130 fungal cells analyzed for WT + 2% FBS, N=127 fungal cells analyzed for WT + 2% human serum, N=120 fungal cells analyzed for GATA1-KO + 2% FBS, N=94 fungal cells analyzed for GATA1-KO + 2% human serum, N=81 fungal cells analyzed for primary neutrophils + 2% FBS, N=101 fungal cells analyzed for primary neutrophils + 2% human serum)(*p<0.05, via a two-way ANOVA with Sidak’s multiple comparison test). (F) Quantification of the fold change in hyphal growth of living fungal cells at 4 hpi relative to 0 hpi following incubation with either iNeutrophils or primary human neutrophils in RPMI media supplemented with 2% FBS or 2% human serum. (N= number of cells over 3 biological replicates)(N=114 fungal cells analyzed for WT + 2% FBS, N=109 fungal cells analyzed for WT + 2% human serum, N=108 fungal cells analyzed for GATA1-KO + 2% FBS, N=64 fungal cells analyzed for GATA1-KO + 2% human serum)(****p<0.0001, via a two-way ANOVA with Sidak’s multiple comparison test)

The ability of neutrophils to actively sense and aggregate around developing hyphae is essential to their antifungal activity [30, 31]. To provide insight into why GATA1-KO iNeutrophils more effectively kill *A. fumigatus* than WT cells, we assessed the ability of this cell line to cluster around developing hyphae following 4 hours of incubation. Overall, the percentage of fungal cells that displayed neutrophil clusters around them was increased in human serum supplemented conditions when compared to FBS supplemented media, but no significant differences were observed when analyzing the number of clusters between WT and GATA1-KO iNeutrophils (Fig. 4E). However, fungal cells that remained at 4hpi were significantly smaller in GATA1-KO iNeutrophil treated samples than the WT treated samples (Fig. 4F). Thus, although both cells aggregate around developing *A. fumigatus* hyphae at similar rates, their ability to control fungal growth within these clusters is improved in GATA1-KO iNeutrophils.

### GATA1-KO iNeutrophils display increased cell surface expression of antifungal PRRs

To determine if the increased antifungal activity of GATA1-KO iNeutrophils is mediated by changes in the surface expression of antifungal receptors, we performed flow analysis of known human fungal pattern recognition receptors. These receptors include the β(1,3)-glucan receptors dectin-1 [32] and complement receptor 3 (CD11b/CD18 heterodimer) [10, 33–35], as well as the toll-like receptors TLR2 and TLR4 [5]. Additionally, Fc-gamma receptor IIA (FcγIIR) plays an important role in the antifungal immune response against opsonized *A. fumigatus* hyphae and was assessed as well [10]. In accordance with previous reports, cell surface marker staining showed that nearly 100% of both the WT and GATA1-KO iNeutrophils expressed the general myeloid marker CD11b (Fig 2A-B), but that the GATA1-KO iNeutrophil population had significantly more cells that also expressed the neutrophil maturation markers CD15 and CD16 (Fig 2C-D) [28]. When analyzing CD11b+ cells, nearly 100% of both WT and GATA1-KO iNeutrophils expressed both CD18 and FcγIIR (Fig 2E-F). However, a significantly higher proportion of GATA1-KO iNeutrophils expressed dectin-1, TLR2 and TLR4 than WT iNeutrophils. Interestingly, further gating on CD15+CD16+ cells showed that GATA1- KO iNeutrophils had comparable expression levels of all PRRs as primary human neutrophils in this mature population, but that WT iNeutrophils still had significantly fewer cells expressing dectin-1 (Fig 2G-H). Collectively, the increased maturity and PRR expression patterns on GATA1-KO iNeutrophils provides a potential mechanism for their improved fungal defense.

**Figure 2:**
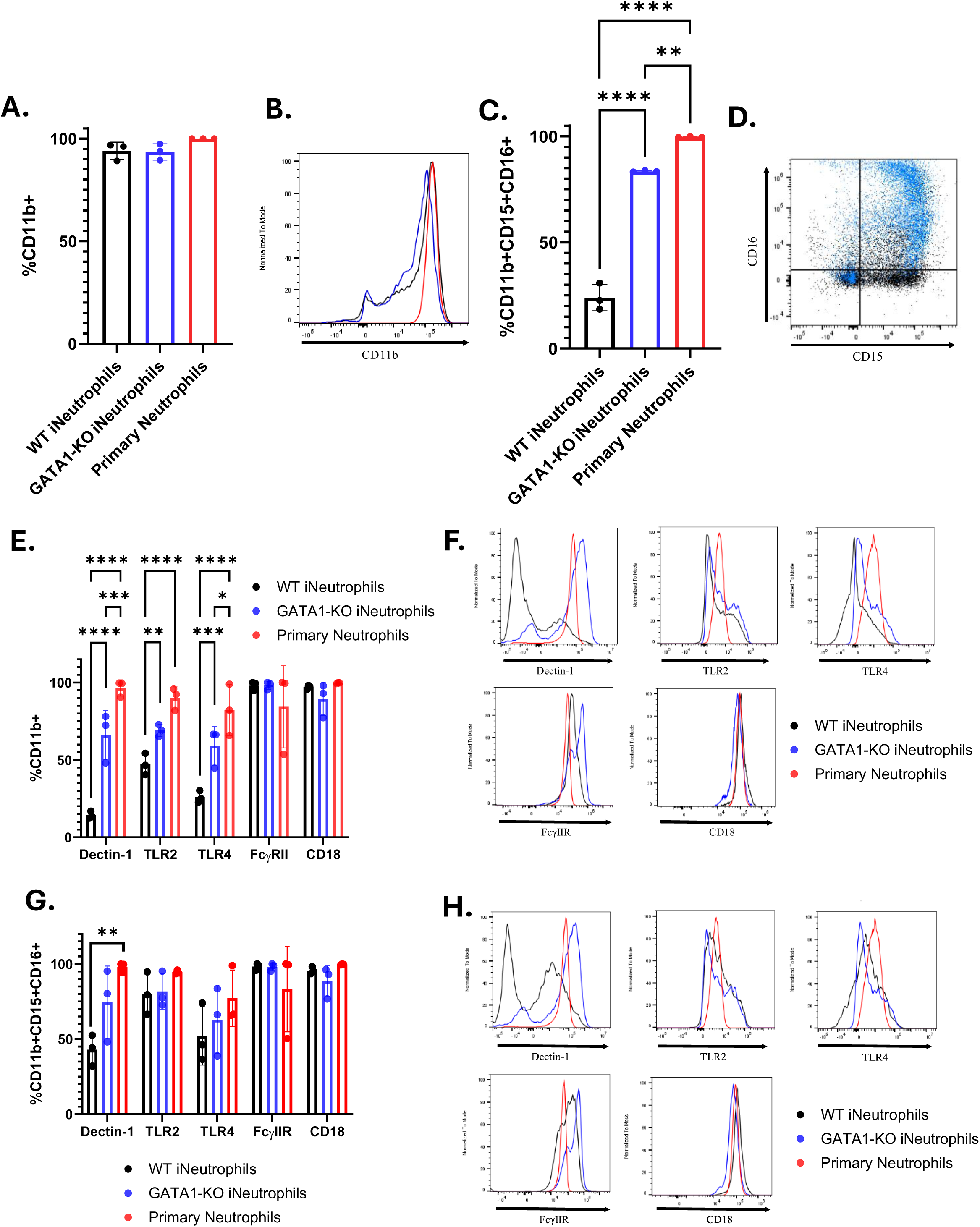
GATA1-KO iNeutrophils display increased surface marker expression of antifungal pattern recognition receptors. (A) Quantification of the percentage of cells expressing CD11b in both iNeutrophils and primary human neutrophils (N=3 biological replicates)(Statistics run via one-way ANOVA with Tukey’s multiple comparison analysis). (B) Representative histogram of CD11b expression intensity on all cell lines stained. Colors of each sample match those shown in 2A. (C) Quantification of the percentage of cells expressing CD11b, CD15 and CD16 in both iNeutrophils and primary human neutrophils (N=3 biological replicates)(**p<0.005, ****p<0.0001, via one-way ANOVA with Tukey’s multiple comparison analysis). (D) Representative scatter plot of CD15 and CD16 expression profiles of WT (black) and GATA1-KO iNeutrophils (blue). (E) Quantification of the percentage of CD11b+ cells expressing antifungal PRRs in both iNeutrophils and primary human neutrophils (N=3 biological replicates)(*p<0.05, **p<0.005, ***p<0.0005, ****p<0.0001, via a two-way ANOVA with Sidak’s multiple comparison test). (F) Representative histograms of PRR expression intensities within the CD11b+ population of all cell lines stained. (G) Quantification of the percentage of CD11b+CD15+CD16+ cells expressing antifungal PRRs in both iNeutrophils and primary human neutrophils (N=3 biological replicates) (*p<0.05, ***p<0.0005, via a two-way ANOVA with Sidak’s multiple comparison test). (H) Representative histograms of PRR expression intensities within the CD11b+CD15+CD16+ population of all cell lines stained.

### GATA1-KO iNeutrophils shift their metabolism toward the pentose phosphate pathway in response to zymosan

PRR binding to microbial pathogens initiates the antifungal activity of neutrophils during infection. Recently, metabolic rewiring towards the pentose cycle following activation in human neutrophils has also been shown to fuel the oxidative burst associated with their antimicrobial defense [14, 15]. To determine if iNeutrophils display a similar shift in metabolism in response to zymosan, a β-glucan rich fungal cell wall bioparticle [36], single cell optical metabolic imaging was performed. This provided single cell autofluorescence lifetime and intensity measurements of reduced nicotinamide adenine dinucleotide (phosphate) (NAD(P)H) and oxidized flavin adenine dinucleotide (FAD) [37]. The findings revealed a decrease in NAD(P)H mean lifetime and a corresponding reduced redox state (indicated by an increase in the redox ratio) of the GATA1-KO iNeutrophils in both zymosan stimulated and unstimulated experimental groups (Fig. 3A-C)(S1 Table). The data suggest that there is likely an overall increase in both free and total NAD(P)H levels within these cells, and thus increased metabolism in the pathways associated with the production of these metabolites, such as glycolysis and the pentose phosphate pathway. To identify what pathways are specifically altered following stimulation, we next performed LC-MS-based analysis of metabolite abundances in both WT and GATA1-KO iNeutrophils. Following zymosan exposure, a significant increase in metabolites associated with the pentose phosphate pathway was observed in both iNeutrophil cell lines exposed to zymosan (Fig 3D). This change also correlated with significant reductions in ATP and fructose-6-phosphate levels. Yet, there were minimal changes between iNeutrophil lines for other metabolites involved in glycolysis and oxidative phosphorylation following stimulation (Fig. 3E-F). Thus, it appears that both WT and GATA1-KO iNeutrophils upregulate pentose phosphate pathway upon activation, similar to what has been reported for primary human neutrophils.

**Figure 3:**
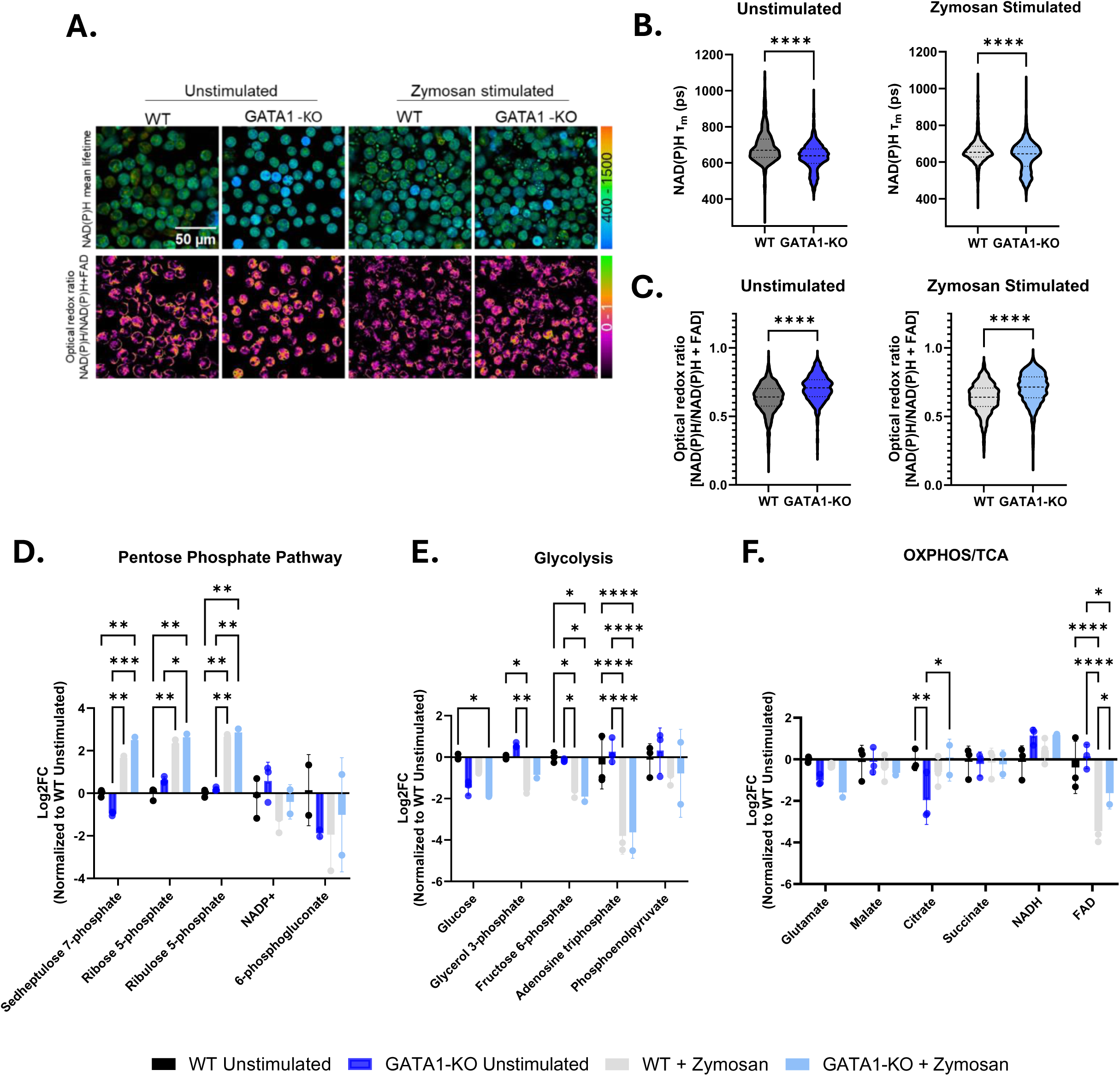
GATA1-KO iNeutrophils rewire their metabolism following activation with zymosan. Representative image of NAD(P)H mean lifetime (top) and optical redox ratio [NAD(P)H/NAD(P)H+FAD] (bottom) used to determine the redox state of WT and GATA1-KO iNeutrophils in unstimulated and zymosan stimulated conditions. Single cell quantification of the (B) NAD(P)H mean lifetime, τ_m_, and (C) optical redox ratio of WT and GATA1-KO iNeutrophils following stimulation with 100μg/ml zymosan. (N=3 biological replicates, 5-6 images per condition per replicate with at least 412 cells per replicate) (****p<0.0001, via one-way ANOVA with Tukey’s multiple comparison posthoc analysis). (D-F) Fold-change changes in the abundances of metabolites associated with the (D) pentose phosphate pathway (E) glycolysis and (F) the oxidative phosphorylation (OXPHOS) pathway and tricarboxcylic acid (TCA) cycle following stimulation with 100μg/ml zymosan. All samples are normalized to the WT unstimulated experimental group. (N=3 biological replicates)(*p<0.05, **p<0.005, ***p<0.0005, ****p<0.0001, via a two-way ANOVA with Sidak’s multiple comparison test).

### GATA1-KO iNeutrophils display enhanced effector functions *in vitro*

To kill fungi, neutrophils generate reactive oxygen species (ROS), release granules and antimicrobial peptides, and form neutrophil extracellular traps (NETs) [6–10]. The ability of these iNeutrophil cell lines to perform these effector functions is critical to their antifungal efficacy. The observation that GATA1-KO iNeutrophils have increased antifungal activity against *A. fumigatus* suggests that they likely have improved effector functions. Indeed, GATA1-KO iNeutrophils have already been reported to produce more NETs than WT cells [28]. However, the response to fungal cell wall epitopes had not been previously examined. Therefore, we exposed iNeutrophils to zymosan and assessed their ability to phagocytose, generate ROS and perform NETosis following activation. GATA1-KO iNeutrophils more efficiently phagocytosed fluorescent pHrodo zymosan than WT iNeutrophils when comparing phagocytosis rates of the CD11b+ populations of both cell lines (Fig. 4A). Similarly, GATA1-KO iNeutrophils showed a shift in ROS production in response to zymosan toward levels found with primary human neutrophils, although this change was not statistically significant when compared to WT cells (Fig. 4B). GATA1-KO iNeutrophils also displayed increased NET formation in response to phorbol myristate acetate (PMA), comparable to primary human neutrophils (Fig. 4C). Collectively, GATA1-KO iNeutrophils show improved effector functions that likely contribute to their increased antifungal activity.

**Figure 4:**
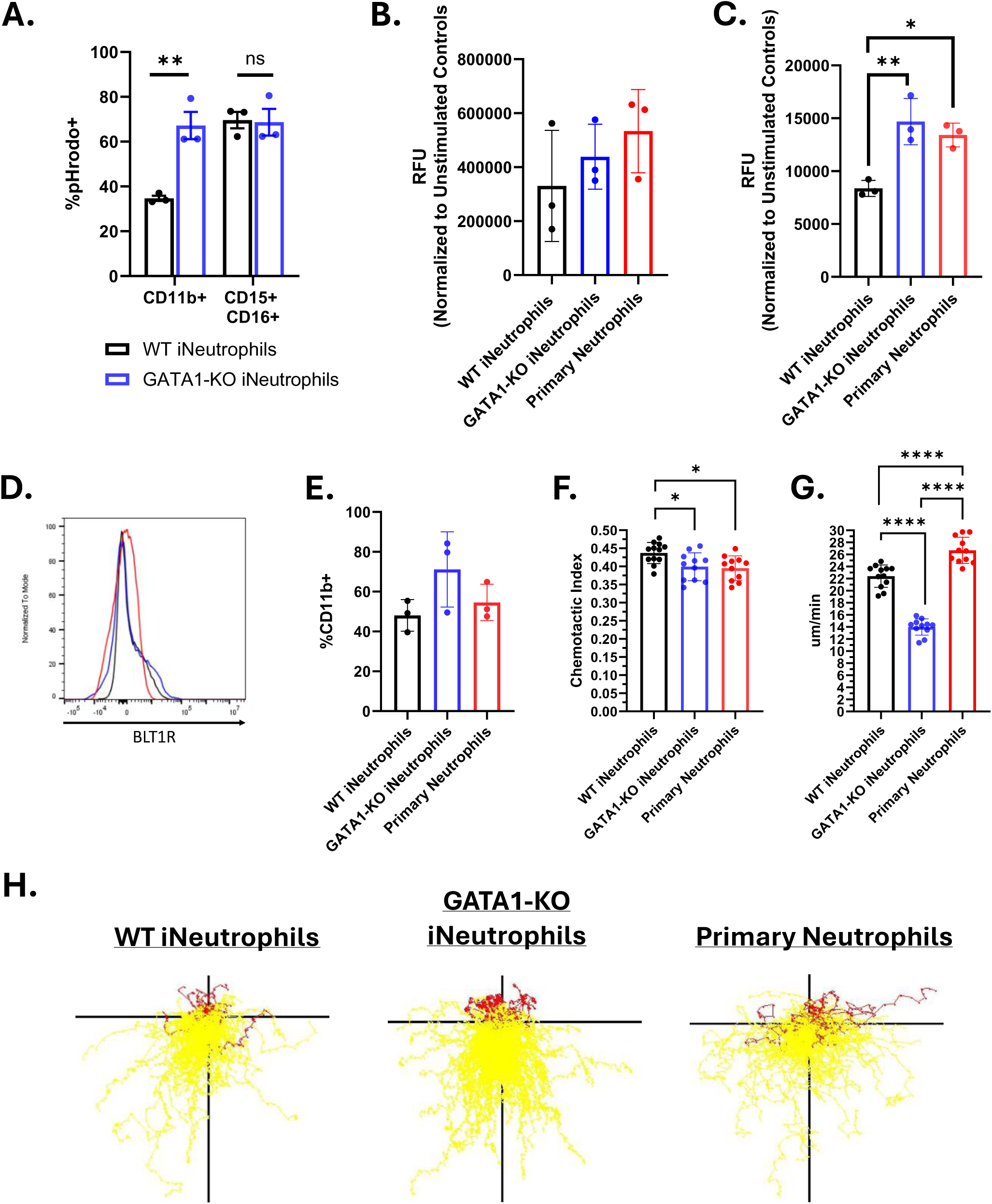
GATA1-KO iNeutrophils effectively migrate and execute effector functions *in vitro*. (A) iNeutrophil phagocytosis of pHrodo zymosan beads quantified by flow cytometry. The percent of CD11b+pHrodo+ and CD11b+CD15+CD16+pHrodo+ cells for both WT and GATA1-KO iNeutrophils are shown. (N=3 biological replicates)(**p<0.005, via a two-way ANOVA with Sidak’s multiple comparison test). (B) Quantification of intracellular ROS production following 2 hours of coincubation with 100μg/ml zymosan. (N=3 biological replicates)(Statistics run via one-way ANOVA with Tukey’s multiple comparison test). (C) Quantification of NET production following 6 hours of coincubation with 50ng/ml PMA. (N=3 biological replicates)(Statistics run via one-way ANOVA with Tukey’s multiple comparison test). (D) Representative histogram of BLT1R expression intensity on both iNeutrophils and primary human neutrophils (colors match those associated with each cell line in figure 4E). (E) Quantification of the number of CD11b+BLT1R+ cells in both iNeutrophils and primary human neutrophils. (N=3 biological replicates)(Statistics run via one-way ANOVA with Tukey’s multiple comparison test). (F) iNeutrophil chemotactic index and (G) mean velocity in response to a LTB4 gradient over 45 minutes of imaging. (N=3 biological replicates with 2-4 technical replicates per test)(*p<0.05, ****p<0.0001, via one-way ANOVA with Tukey’s multiple comparison test) (E) Representative track plots of cell migration through an LTB4 gradient. Yellow tracks indicate forward movement towards higher areas of LTB4 concentration, whereas red tracks indicate cells that moved away.

Central to host defense responses is the ability of neutrophils to migrate and aggregate around invading microbes, and fungal co-incubation has shown that iNeutrophils effectively cluster around growing *A. fumigatus* hyphal structures (Fig. 1). The leukotriene LTB4 is a key player in the process of neutrophil aggregation and swarming in response to fungi [31, 38]. To determine if the iNeutrophil cell lines can similarly respond to LTB4 to mediate their directed migration, we performed cell surface analysis and migration assays. Cell surface marker staining for BLT1R, the high affinity LTB4 receptor [39], showed that both iNeutrophil cell lines expressed the receptor at comparable levels to primary human neutrophils (Fig. 4D-E). Using a previously published microfluidics device and live imaging of neutrophil migration [40], we found that both iNeutrophil cell lines had similar chemotactic responses to LTB4, similar to primary human neutrophils (Fig. 4F-G)(S2 Movie). However, GATA1-KO iNeutrophils had reduced velocity compared to WT cells and primary neutrophils (Fig. 4H). Nonetheless, these results show that GATA1-KO iNeutrophils migrate effectively in response to the chemotactic agent LTB4.

### The integrin CD18 is necessary for *A. fumigatus* killing by GATA1-KO iNeutrophils

Given the enhanced maturity, PRR expression profile and antifungal activity of GATA1-KO iNeutrophils, we propose that this cell line can serve as a model to further characterize neutrophil-fungal interactions *in vitro*. To determine if iNeutrophils provide a genetically tractable model to examine pathways that mediate human neutrophil defense, we tested the effects of deleting a key surface recognition component in neutrophil antifungal responses. Complement receptor 3 (a heterodimeric complex of CD11b/CD18) is the predominant human neutrophil ß(1,3)-glucan receptor that is necessary for fungal recognition and subsequent killing [10, 33–35]. We have shown that GATA1-KO iNeutrophils shift their metabolism following zymosan treatment (Fig. 3), but the role that CR3 has in driving metabolic changes remains unknown. To assess the relationship between CR3-mediated fungal killing and metabolic reprogramming, we deleted CD18 (an integral component of CR3) using CRISPR-Cas9 in GATA1-KO iPSC cells (Fig. S2A-B). Deletion of CD18 did not alter CD15 or CD16 expression levels following iNeutrophil differentiation, suggesting that CD18 does not affect iNeutrophil maturation (Fig. S2C-D). As expected, GATA1-KO/CD18-KO iNeutrophils had reduced levels of CD11b by flow analysis (Fig. S2E-F). In accordance with previous reports in both CD18 deficient human and murine primary neutrophils, the GATA1-KO/CD18-KO double knockout iNeutrophils were almost completely attenuated in their ability to kill *A. fumigatus* germlings over a 10-hour incubation period (Fig. 5A-B) and were unable to control hyphal development over this time (Fig. 5C). Thus, similar to primary neutrophils, GATA1-KO iNeutrophils largely rely on CD18 to mediate their antifungal activity.

**Figure 5:**
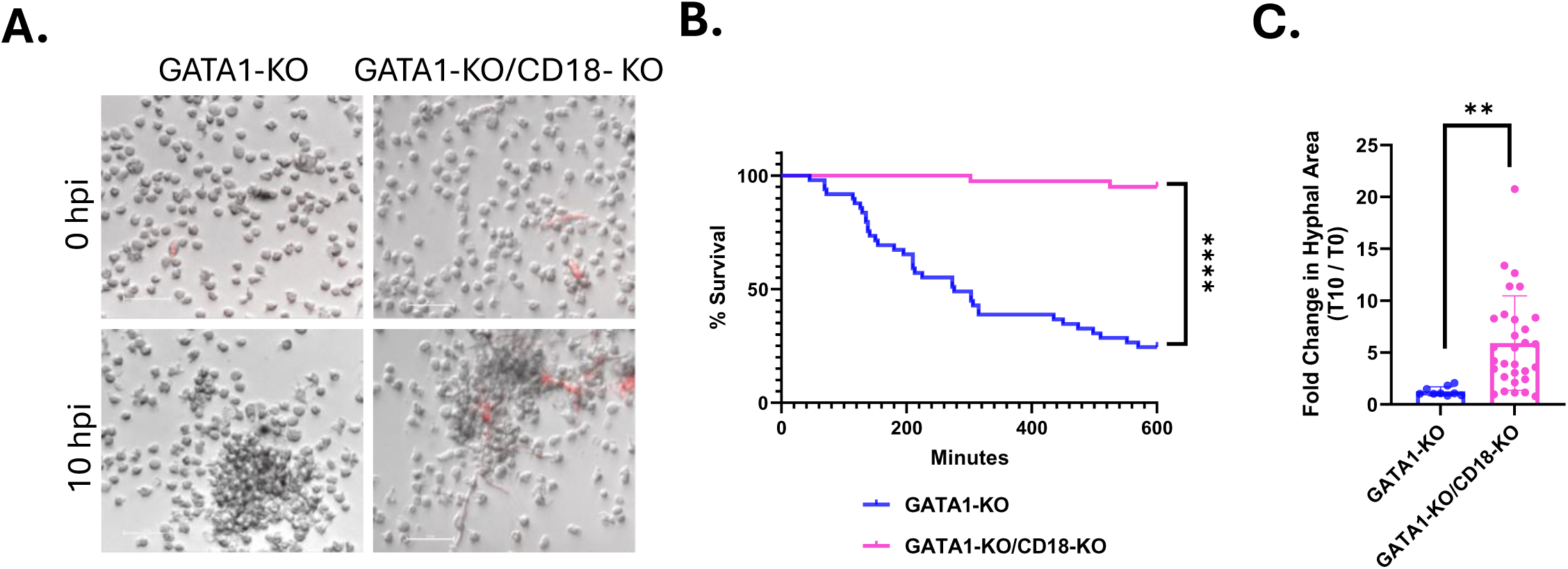
CD18 is necessary for GATA1-KO iNeutrophils to kill and attenuate the growth of *A. fumigatus.* **(A)** Representative image of GATA1-KO and GATA1-KO/CD18-KO iNeutrophils aggregating around and killing RFP expressing *A. fumigatus* germlings following 10 hours of coincubation. (Scale bar = 50μm). (B) Kaplan Meier survival curve of *A. fumigatus* following 10 hours of incubation with either GATA1-KO or GATA1-KO/CD18-KO iNeutrophils in RPMI media supplemented with 2% human serum. Three biological replicates were run for all experimental conditions (N=49 fungal cells tracked for GATA1-KO iNeutrophils and N=40 fungal cells tracked for GATA1-KO/CD18-KO iNeutrophils)(****p<0.0001 via Cox proportional hazard regression analysis). (F) Quantification of the fold change in hyphal growth of living fungal cells at 10 hpi relative to 0 hpi following incubation with either GATA1-KO or GATA1-KO/CD18-KO iNeutrophils. Three biological replicates were run for each condition (N=9 fungal cells assessed for GATA1-KOtreated samples and N=30 fungal cells assessed for GATA1-KO/CD18-KO treated samples)(**p<0.05, via student’s t-test).

### CD18 (CR3) is not required for the upregulation of the pentose phosphate pathway in response to zymosan

CR3 activation has been shown to induce ROS production to aid in the antifungal activity of human neutrophils [35, 41]. Since the GATA1-KO/CD18-KO iNeutrophils are unable to control *A. fumigatus*, we hypothesized that there would be attenuated ROS production following activation. To test this, we incubated both GATA1-KOand GATA1-KO/CD18-KO iNeutrophils with zymosan. Indeed, GATA1-KO/CD18-KO iNeutrophils produced significantly less ROS than the GATA1-KO parental strain following zymosan treatment (Fig. 6A).

**Figure 6:**
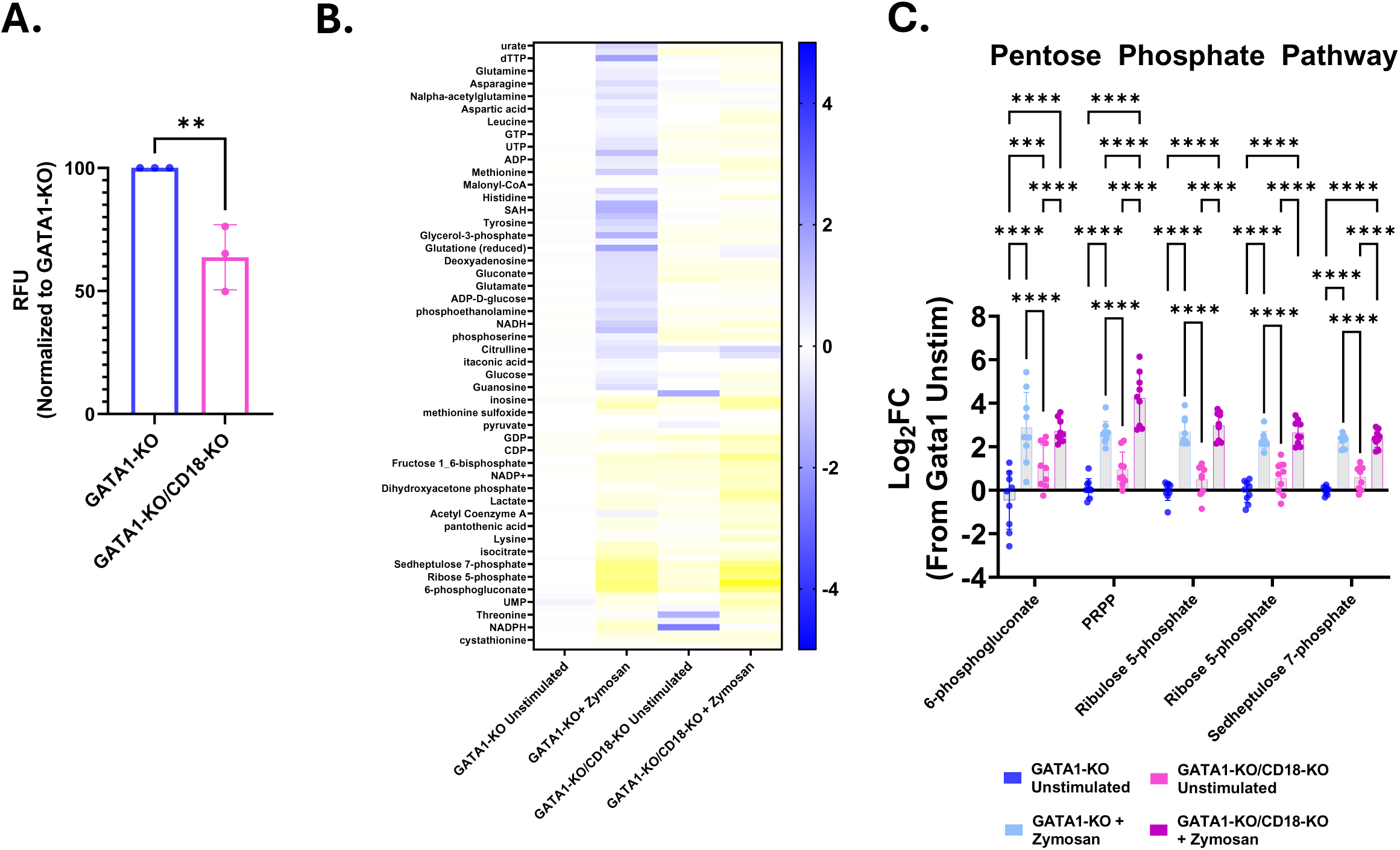
Loss of CD18 (CR3) does not impact metabolic rewiring to the pentose phosphate pathway. (A) Quantification of intracellular ROS production following 2 hours of coincubation with 100μg/ml zymosan. (N=3 biological replicates)(**p<0.05, via student’s t-test). (B) Heatmap showing metabolite abundances in iNeutrophil lines following 2 hours of incubation with 100µg/ml zymosan. Color represents the fold-change compared to GATA1-KO unstimulated control samples. (C) Fold-change in the abundances of metabolites associated with the pentose phosphate pathway. All samples are normalized to the GATA1-KO unstimulated experimental group (N=3 biological replicates with three technical replicates for each test)(***p<0.0005, ****p<0.0001, via a two-way ANOVA with Sidak’s multiple comparison test).

The ability to upregulate the pentose phosphate pathway to enhance ROS production is necessary for efficient antifungal activity [14]. We hypothesized that part of the failure of GATA1- KO/CD18-KO iNeutrophils to kill fungi stems from an inability to shift their metabolism towards the pentose phosphate pathway following activation. Therefore, both GATA1-KO and GATA1-KO/CD18- KO iNeutrophils were stimulated with zymosan and metabolomics was performed. Overall, the metabolic profile of GATA1-KO/CD18-KO iNeutrophils showed differences between GATA1-KO iNeutrophils in both unstimulated and zymosan stimulated conditions (Fig. 6B and S3 Table). However, when specifically assessing metabolites generated from the pentose phosphate pathway, CD18-deficient iNeutrophils shifted their metabolism toward the pentose cycle despite the absence of CR3 (Fig. 6D). Thus, although CD18 is essential to mediate *A. fumigatus* killing, it is not required for upregulation of the pentose phosphate pathway.

### Loss of CD18 impairs the ability of iNeutrophils to efficiently aggregate around developing *A. fumigatus* hyphae

Human neutrophils harboring genetic mutations in *ITGB2* are unable to exit circulation and migrate towards sites of infection [42]. Therefore, we hypothesized that the impaired killing observed in the GATA1-KO/CD18-KO iNeutrophils may be the consequence of migration defects in these cells. To test this, we incubated both GATA1-KO and GATA1-KO/CD18-KO iNeutrophils with *A. fumigatus* germlings and assessed their ability to form neutrophil clusters around developing fungal cells (Fig. 7A). At 2 and 4 hours post incubation, both GATA1-KOand GATA1-KO/CD18-KO iNeutrophil treated samples had comparable numbers of germlings surrounded by neutrophil clusters (Fig. 7B). However, the neutrophil clusters surrounding *A. fumigatus* were significantly smaller with the GATA1-KO/CD18-KO cells (Fig. 7C), suggesting impaired motility and clustering. To further assess this, we performed microfluidic chemotactic assays and found that the CD18-deficient cells had impaired chemotaxis toward LTB4 (Fig. 7D-F and S3 Movie). Taken together, although CD18 is not necessary for fungal-induced upregulation of PPP, it is required for human neutrophil chemotaxis, aggregation and fungal killing.

**Figure 7:**
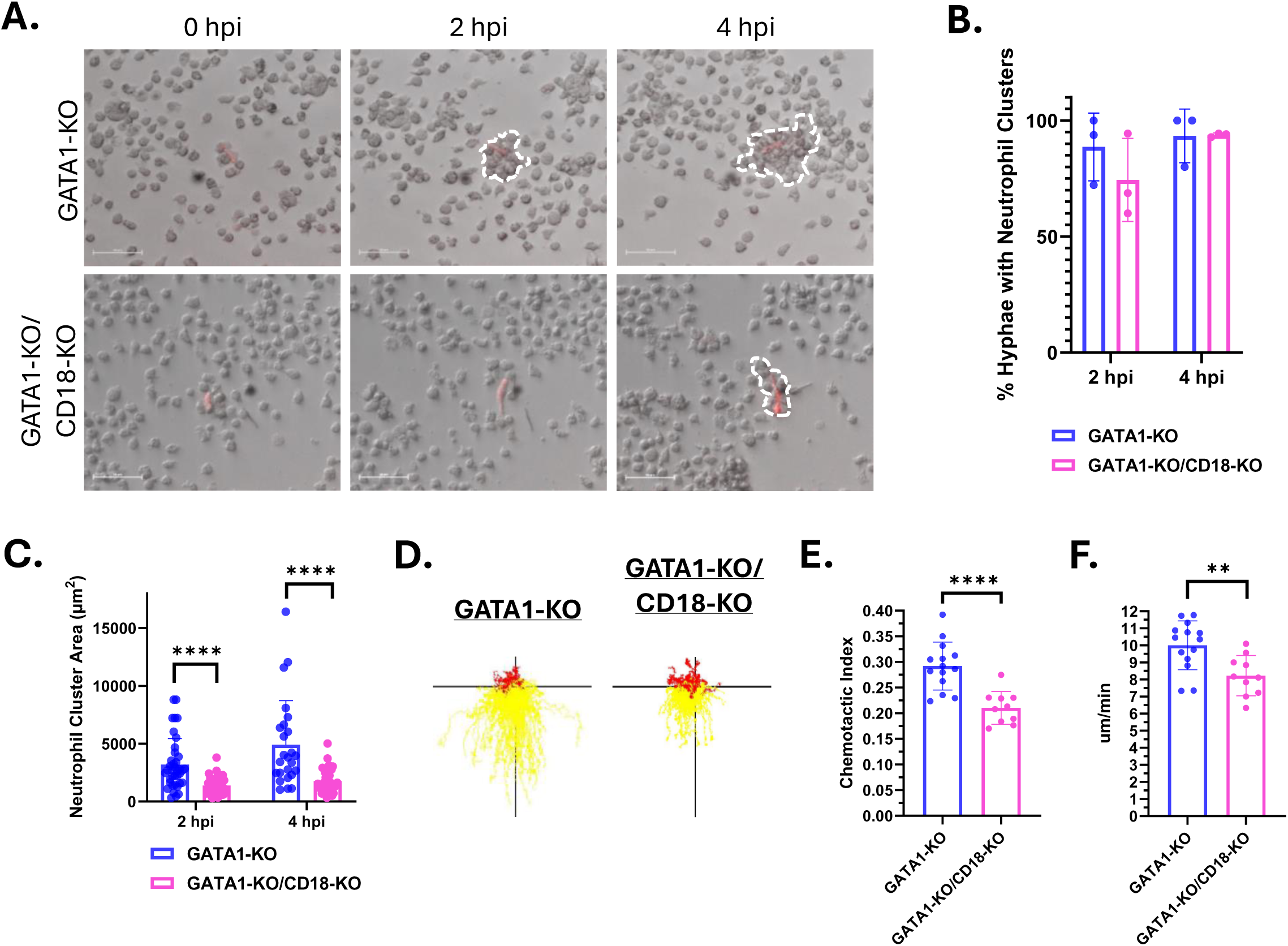
Loss of CD18 impairs the ability of GATA1-KO iNeutrophils to cluster and migrate *in vitro*. (A) Representative time-lapse images of iNeutrophils clustering around *A. fumigatus* hyphae at 2 and 4 hours post incubation. *A. fumigatus* cells are red and iNeutrophil clusters are outlined in dashed white lines. (Scale bar = 50μm) (B) Quantification of the number of fungal cells displaying neutrophil clusters around them at 2 hpi and 4 hpi. Three biological replicates were run for all samples. (N=44 and N=29 fungal cells assessed for GATA1-KO treated samples at 2hpi and 4hpi, respectively. N=57 and N=49 fungal cells assessed for GATA1-KO/CD18-KO treated samples at 2hpi and 4hpi, respectively)(Statistics were run via a two-way ANOVA with a Sidak’s multiple comparison test). (C) Quantification of the size of neutrophil clusters around *A. fumigatus* hyphae at 2 and 4 hours post incubation. (N=38 and N=25 neutrophil clusters assessed for GATA1-KO treated samples at 2hpi and 4hpi, respectively. N=42 and N=41 neutrophil clusters assessed for GATA1-KO/CD18-KO treated samples at 2hpi and 4hpi, respectively)(****p<0.0001, via a two-way ANOVA with a Sidak’s multiple comparison test). (D) Representative track plots of GATA1-KO or GATA1-KO/CD18-KO iNeutrophils migrating through an LTB4 gradient. Yellow tracks indicate forward movement towards higher areas of LTB4 concentration, whereas red tracks indicate cells that moved away. (E) iNeutrophil chemotactic index and (F) mean velocity in response to a LTB4 gradient over 45 minutes of imaging. (N=3 biological replicates with 2-4 technical replicates per test)(**p<0.005, ****p<0.0001, via student’s t-tests).

## Discussion

Here, we show that human iPSC-derived iNeutrophils provide a genetically tractable system to understand human neutrophil antifungal defense. GATA1-KO iNeutrophils are more mature and both attenuate the growth of and directly kill *A. fumigatus* germlings *in vitro* (Fig. 1), thus circumventing previous limitations observed with many existing *in vitro* models. The iNeutrophils also provide a robust system to study the metabolism of human neutrophils and, like primary human neutrophils, shift their metabolism to the pentose phosphate pathway following stimulation. Modification of the iNeutrophils by deletion of the integrin CD18 showed a key role in fungal killing independent of its effects on metabolism. Taken together, our findings support the iNeutrophil system as a powerful tool to understand neutrophil biology.

Deletion of CD18 in iPSCs demonstrated a key role for this integrin in iNeutrophil motility and control of fungal growth. Studies in both humans and mice show that loss of CD18 causes increased susceptibility to bacterial and invasive fungal infections [43, 44]. Additionally, *in vitro* assays using murine neutrophils harboring mutations in CD18 show that this integrin is also needed for phagocytosis, NETosis and ROS production in response to *A. fumigatus* conidia and zymosan [41, 43]. CD18 deficient primary murine neutrophils are also attenuated in their ability to control the growth of *A. fumigatus* hyphae *in vitro* [35, 45]. Here, we show that loss of CD18 in GATA1-KO iNeutrophils recapitulated these phenotypes. Deletion of CD18 almost completely ablated the ability of GATA1-KO iNeutrophils to kill *A. fumigatus* germlings *in vitro*, and significantly impaired their ability to control hyphal growth (Fig. 5). Accordingly, CD18-deficient iNeutrophils were significantly impaired in their migration to fungi and chemoattractants (Fig. 7) and had impaired generation of ROS following stimulation with fungal zymosan (Fig. 6). Importantly, our ability to recapitulate findings from both *in vivo* and *in vitro* phenotypes associated with the loss of CD18 highlights the power of iNeutrophils as an alternative model to study neutrophil antifungal immunity.

Metabolic reprogramming is an important mechanism exhibited by different types of immune cells during host defense to fungi. For example, macrophages, monocytes and natural killer (NK) cells all shift their metabolism towards glycolysis following activation by fungi to mediate their antifungal functions [46–48]. Similarly, glucose transport in neutrophils has been found to be an important mediator of their effector activities in response to *C. albicans*, as loss of the glucose transporter GLUT1 in murine neutrophils attenuated their ability to phagocytose and produce ROS [49]. Recent work has also found that neutrophils further rewire their metabolism towards the pentose phosphate pathway to increase their production of NADPH, and thus increase their capacity to generate ROS [14, 15]. Indeed, this shift has been shown to be important in mediating the antifungal activity of neutrophils against *A. fumigatus in vitro*. To date, only two fungal cell wall ligands have been shown to induce metabolic reprogramming in immune cells, ß-glucan and DHN-melanin [50]. Accordingly, we have shown that iNeutrophils shift their metabolism towards the pentose phosphate pathway following stimulation with ß-glucan rich zymosan (Fig. 3). However, loss of CR3, the primary ß-glucan receptor needed for neutrophil antifungal activity, did not impact upregulation of the pentose phosphate pathway following zymosan exposure (Fig. 6). Collectively, this shows that iNeutrophils provide a powerful tool to study human neutrophil immunometabolism and highlights the need for further research to better understand receptor driven metabolic changes. In summary, we have shown the power of GATA1-KO iNeutrophils as a model to study neutrophil-fungal interaction dynamics *in vitro*. The iNeutrophils also show robust migration to chemoattractants and fungi, reminiscent of a swarming response. These responses are comparable to primary human neutrophils, suggesting that iNeutrophils are an alternative tool to study neutrophil motility in a cell system that are more primary human-like. It is also interesting to consider the potential broader applications of iNeutrophils. Neutropenia leads to increased susceptibility to fungal infections [3]. Granulocyte transfusion therapy has been proposed as a potential means to circumvent these risk factors by providing neutrophils to help combat disease progression [51–53], but limited donor supply and the short lifespan and variable efficacy of primary human neutrophils has thus far limited this approach. The use of human induced pluripotent stem cell (iPSC)-derived neutrophils (iNeutrophils) provides an infinite source of genetically tractable neutrophils as a powerful alternative. Indeed, our protocol produces good manufacturing practice (GMP)-compatible iNeutrophils, opening the possibility for human treatment [26]. The challenge will be to use genetic manipulation to generate iNeutrophils that are optimized for fungal killing but have limited off-target toxicity. These studies will also advance our understanding of how primary human neutrophils are such powerful fungal killing machines.

## Methods

### Fungal growth media and culture conditions

*Aspergillus fumigatus* strain CEA10 expressing a cytosolic RFP protein was used for all fungal experiments (TDGC1.2 - *ΔakuB; argB-; gpdA::RFP::argB; pyrG-; fumipyrG*) [29]. Glycerol stocks were maintained at -80°C and struck out on solid glucose minimal media (GMM) for activation [54]. Once struck out, the plate was incubated at 37°C in the dark for 3 days prior to the experimental start date to induce asexual conidiation. Following incubation, fungal spores were collected in 0.01% Tween 20 solution using an L-spreader and filtered through a 0.4μm mesh filter (Fischer Scientific) into a 50ml conical tube. A 5ml aliquot was then transferred to a new tube, centrifuged at 800rpm for 5 minutes and then resuspended in GMM prior to further back dilution at the start of each experiment.

### Stem cell maintenance and differentiation to iNeutrophils

Differentiation of iNeutrophils from bone marrow-derived human IISH2i-BM9 iPSCs (WiCell) was performed as previously described [55]. Briefly, iPSCs were cultured on Cultrex-coated (WiCell) tissue culture plates in mTeSR-Plus medium (STEMCELL Technologies). To begin the differentiation process, iPSC cells were passaged onto collagen-coated plates (2.4µg/mL) containing TeSR-E8 media with 10µM ROCK inhibitor Y-27632 (ROCKi; Tocris). Cells were then left to adhere to the plate for two hours while incubating at 37°C + 5% CO_2_. Differentiation to hemogenic endothelium (starting at “Day 0”) was initiated via *ETV2* mRNA (TriLink Biotechnologies) transfection in TeSR-E8 media (STEMCELL Technologies) with the use of TransIT reagent and mRNA boost (Mirus Bio). Cells were then left to incubate for one day at 37°C + 5% CO_2_. At day 1, the media was replaced with StemLineII media containing VEGF-165 (20ng/mL; PeproTech) and FGF2 (10ng/mL; PeproTech) to further induce differentiation into hemogenic endothelial cells. This media was then replaced at day 2 with fresh StemLineII media with VEGF-165 and FGF2. At day 3, differentiation from hemogenic endothelia to granulocyte-monocyte progenitors was initiated by changing the media to StemLineII media supplemented with FGF2 (20ng/mL), granulocyte-macrophage colony-stimulating factor (25ng/mL; PeproTech), and UM171 (50nM; Xcess Biosciences). Cells were then left to incubate at 37°C + 5% CO_2_ before topping off the media at day 7 with an equal volume of fresh StemLineII media supplemented FGF2, granulocyte-macrophage colony-stimulating factor and UM171. At day 11, non-adherent progenitor cells were collected and used for iNeutrophil differentiation. Following collection, progenitor cells were cultured in StemSpan SFEM II medium (STEMCELL Technologies), supplemented with GlutaMAX 100× (1×; Thermo Fisher Scientific), ExCyte 0.2% (Merck Millipore), human granulocyte colony-stimulating factor (150ng/mL; PeproTech), and Am580 retinoic acid agonist (2.5μM; STEMCELL Technologies) at 0.8-1 × 10^6^ cells/mL density. After 4 days, suspensions were topped off with equal volumes of StemSpan SFEM II medium supplemented with GlutaMAX, ExCyte, human granulocyte colony-stimulating factor and Am580. iNeutrophils were then collected for experimentation between days 5-7 following initial progenitor harvesting.

### Generation of mutant iPSC cell lines

CRISPR-Cas9 was used as previously described to generate GATA1 and CD18 deficient IISH2i-BM9 iPSCs [23]. To generate a GATA1-KO mutant, two guide RNAs (sgRNAs) were designed with the use of Synthego Guide Design to target exon 2 in the coding sequence of *GATA1* (gRNA1 – 5’ CCAUGGAGUUCCCUGGCCUG 3’ ; gRNA2 – 5’ CAGGAUCCACAAACUGGGGG 3’). Prior to nucleofection, WT iPSC cells were lifted using TrypLE Select (Life Technologies) and singularized by pipetting. Then, 5µg of Cas9 protein (PNA Bio) and 2.5µg of each of the sgRNAs were incubated together for at least 10 minutes at room temperature before nucleofection of iPSC cells occurred using a Human Stem Cell Nucleofector Kit 2 (Lonza). Cells were then plated at 25 cell/cm^2^ on 2x Cultrex-coated plates in mTeSR-Plus media with 1× CloneR supplement (STEMCELL Technologies). Plates were incubated for 2 days before changing the media to mTeSR-Plus. Incubation continued until individual colonies with smooth edges were visible. Then, individual colonies were picked and further expanded in individual wells of a 1x Cultrex-coated 12-well plate. Following expansion, genomic DNA was extracted from individual clones and screened for the loss of a 32 basepair fragment in exon 2 of *GATA1* via polymerase chain reaction (PCR) using forward (5’ ATGGAGACTGAGGTGATGGAGTGG 3’) and reverse (5’ TGCAGCGGTGGCTGTGCTC 3’) screening primers.

To generate a GATA1-KO/CD18-KO double mutant, sgRNAs targeting exon 4 in the coding sequence of *ITGB2* (gRNA1 – 5’ CGAAUGGAGUCAGGAUCCCC 3’ ; gRNA2 – 5’ GCUGCUCAUGAGGGGCUGUG 3’) were designed using Synthego Guide Designer. Nucleofection into the GATA1-KO iPSC parental strain and subsequent expansion of single colonies was then performed as described above. *ITGB2^-/-^* mutants were then confirmed via PCR using the forward (5’ CAGGTCCCGCAGTGTG 3’) and reverse (5’ GTTTCAGCGAGGCTTGTG 3’) screening primers. Mutants were identified by loss of a 52 basepair fragment in the PCR product and confirmed by flow cytometry staining for CD18 following differentiation to iNeutrophils.

### Primary human neutrophil collection

Neutrophils were harvested from blood collected from healthy volunteers under a University of Wisconsin–Madison Minimal Risk Research Institutional Review Board–approved protocol (ID: 2017–0032). Formal written consent was obtained from donors prior to blood draw. Immediately following blood collection, neutrophils were isolated via the use of a MACSxpress negative antibody collection kit (Miltenyi Biotec) per the manufacturer’s instructions. Following isolation, neutrophils were resuspended in 1x phosphate buffered saline (PBS)(Gibco) and subsequently used for different experiments.

### iNeutrophil Fungal Killing Assays

To assess neutrophil/fungal interactions, *A. fumigatus* strain CEA10-RFP (TDGC1.2) was grown as described above. Following spore collection, spores were quantified via hemocytometer and diluted to a final concentration of 4 x 10^3^ cells/ml in GMM. For terminal assays, in which microscopy images were only taken at the start of the experiment and 4 hours post-incubation, 100μl of the spore suspension was added to the appropriate number of wells in a 96-well clear, flat bottom plate previously coated in 10μg/ml fibrinogen and left to incubate at 37°C + 5% CO_2_ for 8 hours, or until germlings appeared. Roughly two hours prior to the end of the spore incubation, primary human neutrophils were collected as previously described and diluted to a cell concentration of 4 x 10^5^ cells/ml in RPMI + 2% heat-inactivated fetal bovine serum (FBS) or 2% pooled human serum. Approximately 20 minutes prior to the end of the spore incubation, iNeutrophils were collected and diluted to a cell concentration of 4 x 10^5^ cells/ml in RPMI + 2% heat-inactivated fetal bovine serum (FBS) or 2% pooled human serum as well. After fungal germling development, the media was removed from all wells in the 96-well plate and replaced with 100μl of the appropriate neutrophil suspension to yield a neutrophil:germling ratio of 100:1. The plate was then immediately taken to a Nikon Eclipse Ti inverted microscope with a preheated chamber set to 37°C and initial images, denoted as time 0 (T0), were taken for all wells. All locations imaged in the plate were tracked in the NIS Element software package connected to the microscope. Following image acquisition, the plate was incubated for 4 hours at 37°C + 5% CO_2_. After 4 hours (T4), the plate was removed and imaged at the same locations captured at the start of the experiment. Images were then analyzed for the presence/absence of red hyphal fluorescence between T4 and T0, and for the formation of neutrophil clusters around hyphal cells that were present at the T4 time point. Additionally, hyphal growth was quantified using Fiji ImageJ (National Institute of Health, Bethesda, MD, USA) by measuring fungal size for all viable fungi at T4 and T0 to determine their fold-change in growth over the incubation period. For all conditions, three biological replicates with three technical replicates were performed using three separate donors for primary human neutrophils or three separate pools of differentiated iNeutrophils. Statistical differences were assessed via a two-way ANOVA with Sidak’s multiple comparison test (GraphPad Prism, v7.0c software).

To further assess the kinetics of fungal killing, time lapse microscopy was performed on (i)neutrophil-fungi cocultures. Following spore collection, 1ml of a 4 x 10^3^ spore suspension in GMM was added to the appropriate number of wells of a 24-well plate previously coated with 10μg/ml fibrinogen. The plate was then left to incubate at 37°C + 5% CO_2_ for 8 hours, or until germlings appeared. During incubation, iNeutrophils and primary human neutrophils were collected and diluted to a concentration of 8 x 10^5^ cells/ml as previously described. Following germling development, GMM media was removed from all wells and replaced with 500μl of the appropriate (i)neutrophil suspension to yield a neutrophil:germling ratio of 100:1. Neutrophil-fungal interactions were then imaged every 3 minutes for 10-12 hours on a Nikon Eclipse TE300 inverted fluorescent microscope (Nikon) with a 20× objective. Environmental controls were set to 37°C with 5% CO_2_. Videos were compiled using the NIS Element software package connected to the microscope and ImageJ software. All experiments were performed using three biological replicates, as defined by three separate donors for primary human neutrophils or three separate pools of differentiated iNeutrophils. Fungal killing was tracked by marking the time point in which fungal RFP signal was lost in all experiments. Statistical tests to measure differences in survival were performed in RStudio using Cox proportional hazard regression analysis with experimental condition included as a group variable, as previously described [56]. Hyphal growth was quantified using Fiji ImageJ by measuring fungal size for all viable fungi at the start of the experiment (T0) and end (T10) to determine their fold-change in growth over the incubation period. Statistical significance was then determined using a student’s t-test (GraphPad Prism, v7.0c software). Neutrophil cluster sizes were measured using Fiji ImageJ at 2 and 4 hours post incubation to assess neutrophil activity as well. Statistical differences were determined via a two-way ANOVA with Sidak’s multiple comparison test (GraphPad Prism, v7.0c software).

### Cell wall staining and flow cytometry

iNeutrophils and Neutrophils were stained in PBS + 1% human serum albumin (HAS) + Brilliant Buffer (Thermo Fisher Scientific) and Human TruStain FcX Fc Receptor Blocking Solution (BioLegend). Following staining, cells were fixed in 2% paraformaldehyde and then analyzed using an Aurora Cytometer (CytekBio). Antibodies used in this study can be found in 1table S4. Data were then analyzed using FlowJo software (v10.8.1; TreeStar). Forward and side scatter parameters were used to differentiate single cells, and Zombie NIR dye was used to assess viability. Within live cells, positive signal for CD11b was used to denote myeloid cells, and mature (i)Neutrophils within that population were further classified by their expression of both CD15 and CD16. The number of cells expressing, and the intensity of their expression profiles, for the antifungal PRRs dectin-1, TLR2, TLR4, CD32 and CD18 within CD11b+ and CD11b+CD15+CD16+ populations were recorded for all cell lines tested.

### Single cell optical metabolic imaging

For optical metabolic imaging, iNeutrophils were plated at 200,000 cells along with the 100μg/mL of zymosan on 35mm glass-bottom dishes (MatTek) about 15 minutes prior to imaging. The dishes were coated with 5µg/ml P-selectin (Bio-Techne) by incubating overnight at 4°C. The cells were housed in a stage top incubator (Tokai Hit) at 37°C and under 5% CO_2_ during the entire duration of imaging.

Optical metabolic imaging was performed on the Ultima Multiphoton Imaging System (Bruker) consisting of an inverted microscope (TI-E, Nikon) coupled to an ultrafast tunable laser source (Insight DS+, Spectra Physics Inc) and time-correlated single-photon counting electronics (SPC -150, Becker & Hickl GmbH) for lifetime measurements. NAD(P)H and FAD were sequentially excited at 750nm and 890nm while their emission was collected using bandpass filters of 460/80nm and 500/100nm, respectively, on GaAsP photomultiplier tubes (H7422P-40, Hamamatsu, Japan). The cells were illuminated using a 40x objective lens (W.I./1.15 NA /Nikon PlanApo). Pixel dwell time of 4.8µs and frame integration of 60 seconds was used with a 2x zoom of field of view (∼0.9 mm^2^). The second harmonic generation (SHG) signal of urea crystals (Sigma-Aldrich) excited at 890nm was collected daily for the instrument response function (IRF).

Fluorescence lifetime data analysis was performed using SPCImage software (Becker & Hickl). To increase photon counts at each pixel, a 3x3 binning containing 9 surrounding pixels was applied. To compute the lifetimes, Weighted Least Squares algorithm was used which entailed an iterative parameter optimization to obtain the lowest sum of the squared differences between model and data. Both NAD(P)H and FAD exist in 2 states depending on their binding status such that NAD(P)H has a shorter lifetime while FAD has a longer lifetime in their free states compared to their respective bound state. Thus the pixel-wise NAD(P)H and FAD decay curves were fit to a biexponential model [I(t) = α_1_ × exp (−τ/ τ_1_) + α_2_× exp(−τ/τ_2_) + C)] convolved with the system IRF such that I(t) is the fluorescence intensity measured at time t, α_1_, α_2_ are the fractional contributions and τ_1_, τ_2_ denote the short and long lifetime components, respectively. C accounts for background light. The goodness of the fit was checked using a reduced chi-squared value<1.0. The mean lifetime is the weighted average of the free and bound lifetimes (τ_m_ = α_1_ × τ_1_ + α_1_ × τ_2_) and is calculated for each pixel. The area under the fluorescence decay curve at each pixel was integrated to generate the intensity of NAD(P)H and FAD. The optical redox ratio at each pixel is computed as the intensity ratio [NAD(P)H / (NAD(P)H + FAD)]. For generating single cell data, single cell masks were computed from the NAD(P)H intensity image using Cellpose 2.0 (model = cyto 2, radius = 30) [57] followed by manual checks and edits on napari [58] . Pixels within cell masks that contained zymosan were manually identified by their shape and size and were excluded from analysis. A custom python library, Cell Analysis Tools [59] ,was used for processing and extracting the single cell data.

### Metabolomic analysis

To assess metabolite abundances following zymosan exposure, 2 x 10^6^ iNeutrophils were collected and transferred to a 1.5ml microcentrifuge tube. Cells were then washed once with PBS and resuspended in 500μl of StemSpan SFEM II medium (STEMCELL Technologies), supplemented with GlutaMAX 100x, ExCyte 0.2%, human granulocyte colony-stimulating factor (150 ng/mL), and Am580 retinoic acid agonist (2.5 μM). Following resuspension, 500μl of zymosan (Invivogen) supplemented media was added to the appropriate tubes to make a final working concentration of 100μg/ml. Similarly, media without zymosan was added to designated tubes to serve as an unstimulated control. Tubes were then left to incubate at 37°C + 5% CO_2_ for two hours, with gentle mixing via pipetting occurring every 40 minutes throughout the incubation process. Following incubation, cells were centrifuged at 200xg for 5 minutes, the supernatant was removed and the cell pellet subsequently washed with 1ml of PBS. After washing, 500μl of an ice cold 80:20 methanol:water (MeOH:H_2_O) solution was added to each tube, vortexed and then placed in a -80°C freezer for 30 minutes. Following incubation, tubes were vortexed again and spun at 13,000rpm for 5 minutes at 4°C. The supernatant was transferred to a new microcentrifuge tube over dry ice. 150μl of an ice cold 80:20 MeOH:H_2_O solution was then added to the tube containing the original pellet, and the tube was vortexed, centrifuged and the supernatant was transferred to the same tube corresponding to each sample as was done before. For each condition tested, extraction of three biological replicates with three technical replicates each was performed for analysis.

Metabolites were measured using a Thermo Q-Exactive mass spectrometer coupled to a Vanquish Horizon UHPLC. Data were collected with Xcalibur 4.0 software (Thermo) and peak integration was performed using Maven [60, 61]. The data was collected on a full scan negative mode. The metabolites identified were based on exact m/z and retention times that were determined with chemical standards. After isolation in -80°C 80:20 MeOH:H_2_O, metabolite extracts were dried under nitrogen stream. Samples were resuspended in LC-MS grade water and separated on a 2.1 × 100mm, 1.7μM Acquity UPLC BEH C18 Column (Waters). The solvents used were A: 97:3 H_2_O:MeOH (v:v), 10mM tributylamine, 9 mM acetate, pH 8.2 and B: 100% MeOH. The gradient was 0 min, 95% A; 2.5 min, 95% A; 17 min, 5% A; 21 min, 5% A; 21.5 min 95% A. The flow rate was 0.2 ml/min and the column temperature was 30°C. Setting for the ion source were; 10 aux gas flow rate, 35 sheath gas flow rate, 2 sweep gas flow rate, 3.2kV spray voltage, 320°C capillary temperature and 300°C heater temperature.

### Phagocytosis assays

The phagocytic capabilities of (i)Neutrophils were assessed with the use of pHrodo Green zymosan BioParticles (Invitrogen) via the manufacturer’s instructions. Briefly, the BioParticles were opsonized with 30% pooled human serum (#MP092930149; MP Biomedicals) for 30 minutes at 37°C then washed 3 times in PBS. One million cells were resuspended in 80µL of StemSpan SFEM II medium, then20 µL of opsonized beads were added, giving a 100:1 multiplicity of infection. Cells and beads were incubated in an Eppendorf tube for 1 hour at 37 °C, then stopped by addition of ice-cold PBS. While keeping tubes on ice, cells were stained with neutrophil lineage markers (Supplementary Table 1), then fixed with 2% PFA before flow cytometry analysis on the Aurora Cytometer (CytekBio).

### Reactive oxygen species quantification

Intracellular peroxynitrate production was assessed via DHR123 (Invitrogen) staining. Briefly, 100μl of 1 x 10^5^ cells of either iNeutrophils or primary human neutrophils in RPMI + 2% FBS + 5μg/mL DHR123 was added to wells of a black 96-well clear-bottom plate previously coated with 10μg/mL fibrinogen (Sigma). Cells were then left to incubate for 1 hour at 37°C + 5% CO_2_ before a working concentration of 100μg/mL of zymosan was added to the appropriate wells. Control wells containing a corresponding volume of media without zymosan were generated as well. Following zymosan addition, plates were incubated in the dark at 37°C + 5% CO_2_ before their fluorescent signal was measured at 2 hours post-incubation at 485/535nm using the Victor3V microplate reader (PerkinElmer). Background signal from unstimulated cells were subtracted from zymosan stimulated samples for analysis. Three biological replicates were run for all cell lines tested, and statistical analysis was performed using either a student’s t-test or a one-way ANOVA with a Tukey’s multiple comparison analysis (GraphPad Prism, v7.0c software).

### NETosis assays

NET release was quantified by measuring extracellular DNA via SYTOX green staining as previously described [29]. To achieve this, 2 x 10^5^ cells in RPMI + 2% FBS media were added to wells of a black, clear-bottom 96-well plate previously coated with 10μg/mL fibrinogen. PMA was then added to a working concentration of 100nM to the appropriate wells. Control wells containing a corresponding volume of media without PMA were generated as well. The plate was then left to incubate in the dark at 37°C + 5% CO_2_ for 6 hours before adding SYTOX Green (Thermofisher) to a working concentration on 375nM to all wells. Fluorescence (485nm/535nm, 0.1s) was measured 15 minutes after addition of SYTOX Green using a Victor3V plate reader (PerkinElmer). Background signal from unstimulated cells were subtracted from PMA stimulated samples for analysis. Three biological replicates were run for all cell lines tested, and statistical analysis was performed using a one-way ANOVA with a Tukey’s multiple comparison analysis (GraphPad Prism, v7.0c software).

### PrestoBlue staining for fungal viability

PrestoBlue staining was used to assess fungal growth following coincubation with (i)neutrophils. Following collection, spores were quantified via hemocytometer and diluted to a final concentration of 4 x 10^3^ cells/ml in GMM. 100μl of the spore suspension was then added to the appropriate number of wells in a 96-well clear, flat bottom plate previously coated in 10μg/ml fibrinogen and left to incubate at 37°C + 5% CO_2_ for 8 hours, or until germlings appeared. Following germling appearance, the media was removed and 100μl of a 4 x 10^5^ cells/ml solution of either iNeutrophils or primary human neutrophils in RPMI + 2% FBS or 2% human serum were added to each well. Additionally, wells containing growth media with fungus alone were created to serve as a control to quantify fungal attenuation. All wells were then topped off with an additional 100μl of the appropriate media to prevent them from drying out during incubation and then plates were left for 24 hours at 37°C + 5% CO_2_. Once the incubation was complete, the plate was centrifuged at 3500rpm for 5 minutes to pellet cellular material to the bottom of the plate. The media was then gently removed, replaced with 200ul of a neutrophil lysis solution (basic H_2_O (pH 11) + 100 100μg/ml DNAse I) and then left to incubate for 30 minutes at 37°C. This process was then repeated two additional times or until (i)neutrophils were visibly lysed under brightfield microscopy. Following (i)neutrophil lysis, the media was replaced with GMM + PrestoBlue viability reagent (1:10 dilution, ThermoFisher) and the plate was left to incubate for 3 hours at 37°C. Following incubation, media was removed and added to fresh wells that did not contain fungi and fluorescence was measured at 555/590nm using a PerkinElmer Victor3V plate reader. Three biological replicates were run for all cell lines tested in each condition, and statistical analysis was performed using a two-way ANOVA with a Sidak’s multiple comparison test (GraphPad Prism, v7.0c software).

### LTB4 Chemotaxis Assays

Chemotaxis was assessed using a microfluidic device as described previously [40]. In brief, polydimethylsiloxane (PDMS) devices were plasma treated and adhered to glass coverslips. Devices were coated with 10μg/mL fibrinogen + 1ug/ml fibronecin + 1.2ug/ml Collogen IV (Sigma) in PBS for 30 min at 37°C + 5% CO_2_ or at 4°C overnight. The devices were then blocked with 2% bovine serum albumin (BSA)-PBS for 30 min at 37°C + 5% CO_2_, and then washed twice with PBS. Cells were stained with Calcein AM (Molecular Probes) in PBS for 20 min at 37°C + 5% CO_2_, washed once with PBS and then resuspended in imaging media (0.25% StemSpan-II, 0.5% FBS in PBS). Cells were seeded at 8 × 10^6^ cells/mL and allowed to settle for 10 minutes before addition of chemoattractant. 3μl of 3μM LTB_4_was loaded into the input port of the microfluidic device. Cells were imaged every 30 seconds for 60 minutes on a Nikon Eclipse TE300 inverted fluorescent microscope with a 10× objective and an automated stage using MetaMorph software (Molecular Devices). Automated cell tracking analysis was done using JEX software [62] to calculate chemotactic index and velocity and to generate representative rose plots for migration patterns.

## Acknowledgments

We would like to thank the members of the Huttenlocher and Keller labs for their important insights and discussions of the research throughout the development of this manuscript.

## Funding

This work was supported by R21AI184357 awarded to A.H. from the National Institute of Allergy and Infectious Diseases (NIAID) of the National Institutes of Health (NIH). ASW was supported by F32AI183696-01 from the NIAID (NIH) and T32ES007015 of the NIH.

## Competing Interests

The authors declare that no competing interests exist.

## Author Contribution

**Conceptualization:** Andrew S. Wagner, Frances M. Smith, Nancy Keller and Anna Huttenlocher

**Data Curation:** Andrew S. Wagner, James A. Votave, Rupsa Datta

**Formal Analysis:** Andrew S. Wagner, Frances M. Smith, David A. Bennin, James A. Votava, Rupsa Datta, Morgan A. Giese and Wenxuan Zhao

**Funding Acquisition:** Andrew S. Wagner, Melissa C. Skala, Jing Fan, Nancy P. Keller and Anna Huttenlocher

**Investigation:** Andrew S. Wagner, Frances M. Smith, David A. Bennin, James A. Votava, Rupsa Datta, Morgan A. Giese and Wenxuan Zhao

**Methodology:** Andrew S. Wagner, Frances M. Smith, David A. Bennin, James A. Votava, Rupsa Datta and Morgan A. Giese

**Project Administration:** Nancy Keller and Anna Huttenlocher

**Resources:** Melissa C. Skala, Jing Fan, Nancy P. Keller and Anna Huttenlocher

**Software:** James Votava, Rupsa Datta and Wenxuan Zhao

**Validation:** Andrew S. Wagner

**Visualization:** Andrew S. Wagner

**Writing - Original Draft:** Andrew S. Wagner

**Writing - Review and Editing:** Frances M. Smith, David A. Bennin, James A. Votava, Rupsa Datta, Morgan A. Giese, Wenxuan Zhao, Melissa C. Skala, Jing Fan, Nancy P. Keller and Anna Huttenlocher

---

**S1 Fig:**
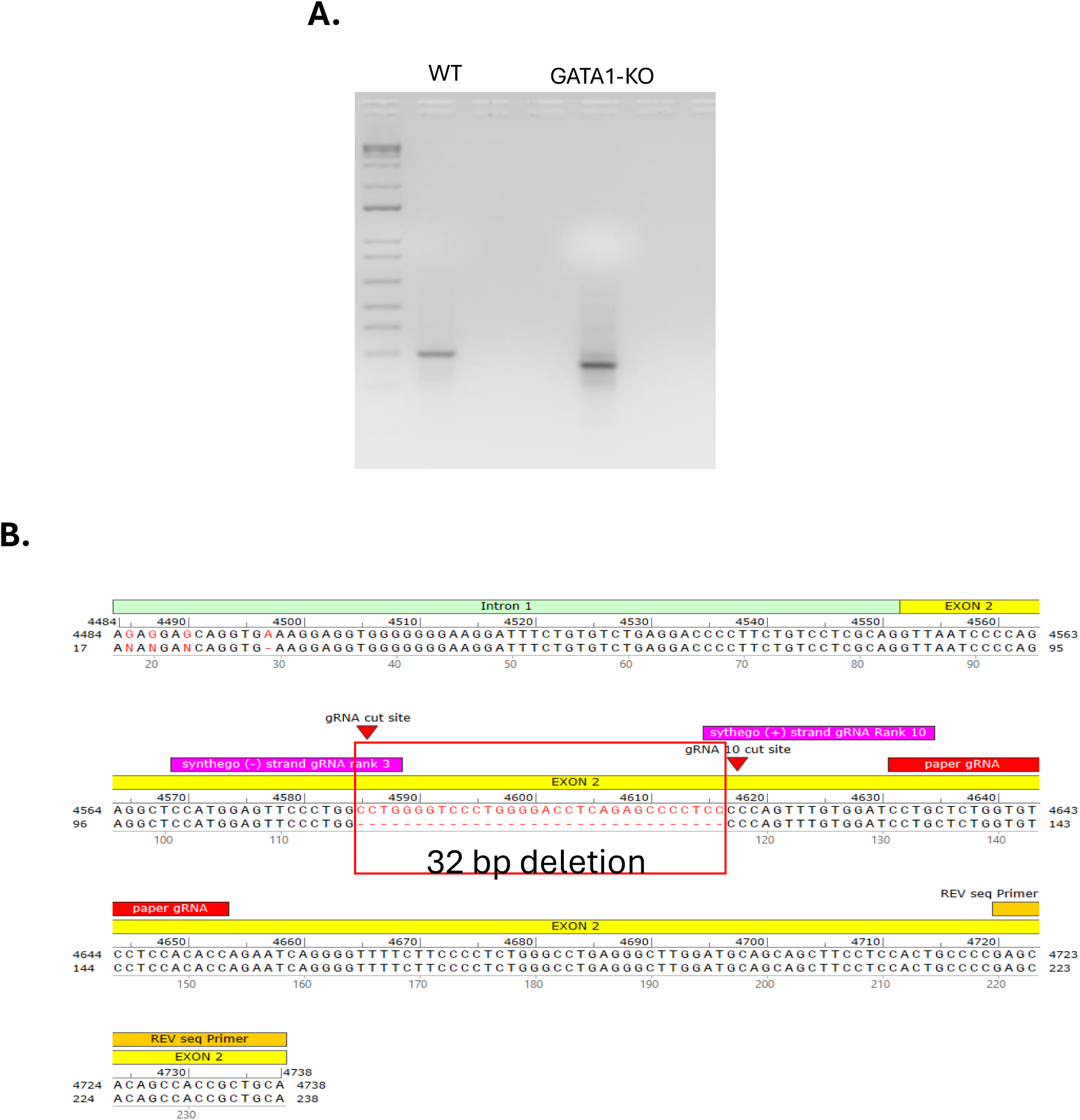
Confirmation of 32bp deletion in the ORF of *GATA1*. (A) Agarose gel showing a shift in the *GATA1* gene following CRISPR-Cas9 mediated deletion of a 32bp fragment in exon 2 of the coding sequence for the gene. (B) Sanger sequencing results confirming loss of the 32bp fragment in exon 2 of the GATA1-KO mutant.

**S2 Fig:**
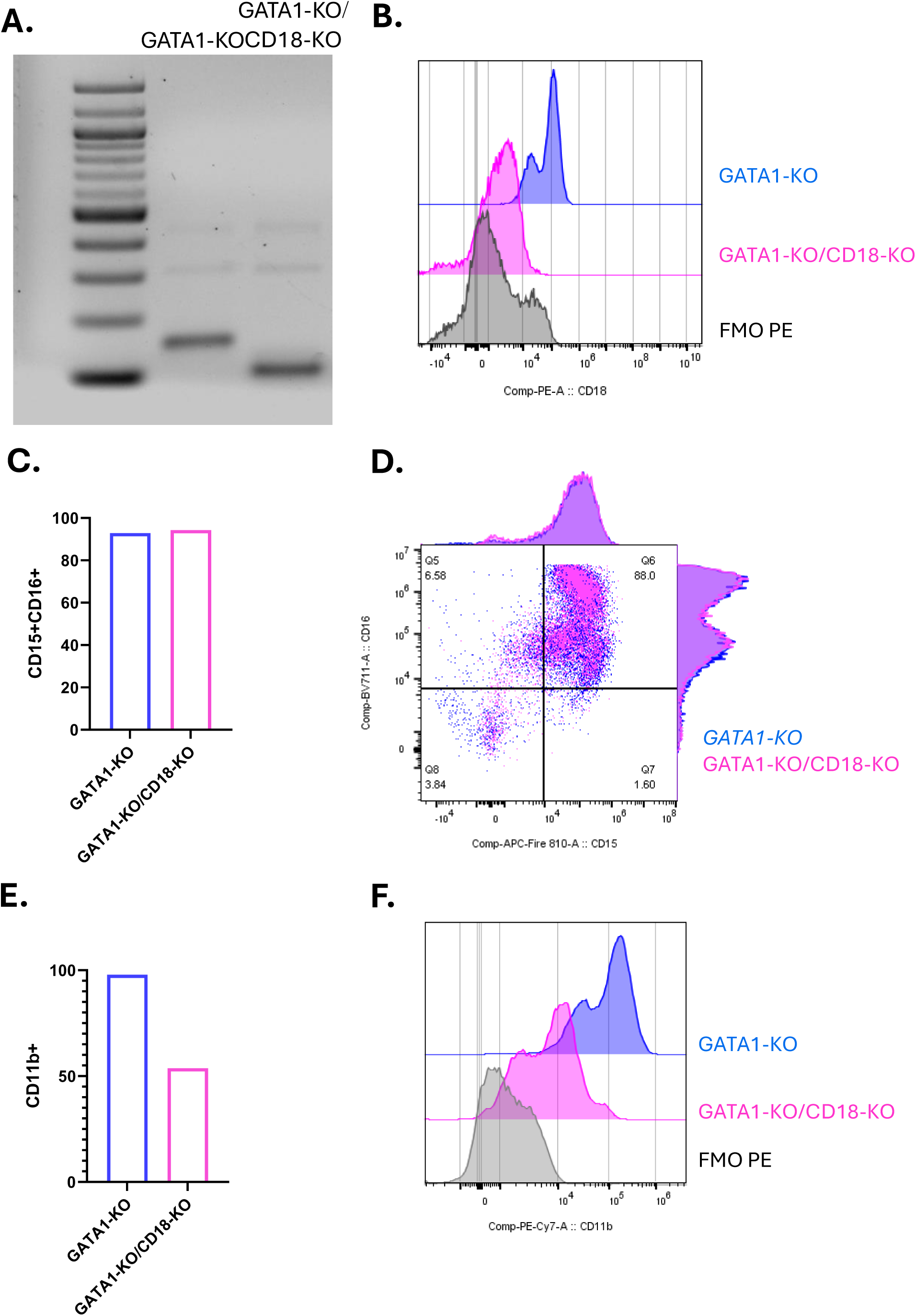
Loss of CD18 reduces surface expression of CD11b. (A) Agarose gel showing a shift in the *ITGB2* gene following CRISPR-Cas9 mediated deletion of a 52bp fragment in exon 4 of the coding sequence of the gene. (B) Representative histograms of surface marker staining for CD18 in GATA1-KO and GATA1-KO/CD18-KO iNeutrophils. Fluorescence minus one (FMO) PE is indicative of a negative control for CD18 staining where the anti-CD18-PE antibody was not added to the cells. (C) Quantification of the number of CD15+CD16+ cells within the live population of iNeutrophils. (D) Representative scatter plots of CD15 and CD16 expression intensities for GATA1-KO (blue) and GATA1-KO/CD18-KO (pink) iNeutrophils. (E) Quantification of CD11b+ cells within the live population of iNeutrophils. (F) Representative histograms of CD11b expression intensities.

**S1 Movie: GATA1-KO iNeutrophils aggregate around and kill *A. fumigatus in vitro*.** Representative movie of GATA1-KO iNeutrophils interacting with and killing *A.fumigatus* germlings (red) during coincubation. Images were taken every 3 minutes over a 12-hour incubation period. Scale bar is 50µm. Movie is displayed as 7 frames/second.

**S2 Movie: iNeutrophils migrate efficiently in response to the chemoattractant LTB4.** Representative fluorescent movies of WT (left) and GATA1-KO (center) and primary human (right) (i)Neutrophils migrating towards an LTB4 gradient at the bottom of the image. iNeutrophils are stained with calcein and tracked using the fluorescent images. Cell tracks of quantified cells are overlaid and appear yellow. Scale bar is 100µm. Images were taken every 30 seconds for 45 minutes. Movie is displayed as 7 frames/second.

**S2 Movie: Loss of CD18 impairs GATA1-KO iNeutrophil migration efficiently to the LTB4.** Representative fluorescent movies of GATA1-KO (left) and GATA1-KO/CD18-KO (right) iNeutrophils migrating towards an LTB4 gradient at the bottom of the image. iNeutrophils are stained with calcein and tracked using the fluorescent images. Cell tracks of quantified cells are overlaid and appear yellow. Note the reduced track length of GATA1-KO/CD18-KO cells compared to their parental control. Scale bar is 100µm. Images were taken every 30 seconds for 45 minutes. Movie is displayed as 7 frames/second.

**Supplementary Table 1:** Antibodies used in this study.

| Target | Fluor | Clone | Catalog # | Vendor |
| --- | --- | --- | --- | --- |
| CD11b | PeCy7 | ICRF44 | 301321 | Biolegend |
| CD15 | APC-Fire 810 | W6D3 | 323058 | Biolegend |
| CD16 | BV711 | 3G8 | 563127 | BD Biosciences |
| BLT1R | BUV805 | 14F11 | 749044 | BD Biosciences |
| Clec7a (Dectin-1) | APC | 15E2 | 355405 | Biolegend |
| TLR2 | FITC | W15145C | 392307 | Biolegend |
| TLR4 | PE | HTA125 | 312805 | Biolegend |
| CD18 | PE | CBRLFA-1/2 | 366304 | Biolegend |
| CD32 | FITC | FUN-2 | 303204 | Biolegend |
| Zombie NIR | 746 | NA | 423105 | FischerScientific |
| Human TruStain<br>FcX Fc Receptor<br>Blocking Solution | NA | NA | 422302 | Biolegend |
| Ultracomp eBeads | NA | NA | 501129040 | ThermoFisher |

## Notes

### Competing Interest Statement

The authors have declared no competing interest.

